# Role of Innate-like Lymphocytes in the Pathogenesis of Community Acquired Pneumonia

**DOI:** 10.1101/495556

**Authors:** RF Hannaway, X Wang, M Schneider, S Slow, MR Schofield, X Morgan, DR Murdoch, JE Ussher

## Abstract

**Background:** Mucosal-associated invariant T (MAIT) cells and Vδ2^+^ γδ T cells are anti-bacterial innate-like lymphocytes (ILLs) that are enriched in blood and mucosa. ILLs have been implicated in control of bacterial infection. However, the role of ILLs in community-acquired pneumonia (CAP) is unknown.

**Methods:** Using sputum samples from a well-characterised CAP cohort, MAIT cell (Vα7.2-Jα12/20/33) and Vδ2^+^ T cell (Vδ2-Jδ1/2/3/4) abundance was determined by quantitative PCR. Cytokine and chemokine concentrations in sputum were measured. The capacity of bacteria in sputum to produce activating ligands for MAIT cells and Vδ2^+^ T cells was inferred by 16S rRNA sequencing.

**Results:** MAIT cell abundance in sputum was higher in patients with less severe pneumonia; duration of hospital admission was inversely correlated with both MAIT and Vδ2^+^ T cell abundance. The abundance of both ILLs was higher in patients with a confirmed bacterial aetiology, however there was no correlation with total bacterial load or the predicted capacity of bacteria to produce activating ligands. Sputum MAIT cell abundance was associated with interferon- α, and interferon-*γ*, and sputum neutrophil abundance, while Vδ2^+^ T cell abundance was associated with CXCL11 and interferon-*γ*.

**Conclusions:** Pulmonary MAIT and Vδ2^+^ T cells can be detected in sputum in CAP, where they may contribute to improved clinical outcome.

## Introduction

Community-acquired pneumonia (CAP) is a leading cause of hospital admission [1], Innate immune defences are critical for the prevention and early control of pulmonary infection, and for instructing the adaptive immune response [2]. Innate and adaptive immunity are bridged by innate-like lymphoid cells, such as mucosal associated invariant T (MAIT) cells and γδ T cells; however, their role in CAP is not yet fully understood.

MAIT cells occur at high frequencies in blood and at mucosal surfaces, including the lung [3]. They have a semi-invariant T cell receptor (TCR), Vα7.2-Jα12/20/33, that is restricted by the highly-conserved, MHC class Ib-like molecule, MR1, and recognises derivatives of the riboflavin synthesis pathway [4, 5]. This pathway is present in many bacteria, including numerous pulmonary pathogens, but not in humans [6]. In addition, MAIT cells can be activated independently of their TCR by cytokines, in particular, by interleukin-12 and -18, suggesting potential involvement in the immune response to non-riboflavin-synthesising bacteria and viruses [7]. Studies in mice suggest that MAIT cells play an important role in the protection against both bacterial and viral pulmonary pathogens and in the containment of chronic infection [8–11]. In humans, several observations suggest a potential role of MAIT cells in protection against pneumonia. Firstly, MAIT cell frequency in blood is decreased in patients with pneumonia or active tuberculosis [12]. Secondly, in critically ill patients, a persistently low frequency of MAIT cells increases the risk of subsequent nosocomial infections, including pneumonia [13]. Thirdly, MAIT cells are depleted in HIV-infected patients [14], who are 25 times more likely to develop bacterial pneumonia, an increased risk which is not completely explained by the loss of CD4^+^ T cells [15].

The Vγ9Vδ2 subset is the predominant γδ T cell population in humans, comprising 1-9% of circulating T cells. They are unique to humans and higher primates and are activated by (E)-4-hydroxy-3-methyl-but-2-enyl pyrophosphate (HMB-PP), an intermediate of the non-mevalonate pathway of isoprenoid biosynthesis present in many bacteria, including pulmonary pathogens, but not in humans [16]. Vγ9Vδ2 T cells may also play a role in the immune response to pulmonary infection. Expansion of the human Vγ9Vδ2 T cell population has been reported in several bacterial infections [16, 17], as well as their rapid trafficking to the lungs after activation [18]. Adoptive transfer of Vγ9Vδ2 T cells in human primates protected them from pulmonary *M. tuberculosis* infection [19].

In this study, we developed quantitative real-time polymerase chain reaction (qPCR) assays to investigate the abundance of MAIT cells and Vγ9Vδ2 T cells in the sputum of patients with CAP. The abundance of MAIT cells and Vγ9Vδ2 T cells in sputum was correlated with clinical data.

## Methods

### Sputum and blood samples

Frozen (-80°C) sputum samples (n=88) from a previously reported cohort of patients with CAP were analysed [20]. Ethical approval was granted by the Northern A Health and Disability Ethics Committee (12/NTA/30).

To validate the qPCR method, blood from healthy adult volunteers was used (n=13). Peripheral blood mononuclear cells (PBMCs) were isolated using Lymphoprep (Alere Technologies AS, Oslo, Norway) and cryopreserved at -80°C until use. Collection of blood was approved by the University of Otago Ethics Committee (Health) (H14/046).

### Real-time Polymerase Chain Reaction (qPCR)

DNA was extracted from sputum and PBMCs using the NucleoSpin Tissue DNA Kit (Macherey-Nagel, Düren, Germany). Prior to DNA extraction, samples were defrosted and pretreated with dithiothreitol (0.1mg/mL) at 37°C for 20 minutes, or until fully digested. The primers and probes used for quantification of MAIT cells (Vα7.2-Jα12/20/33), Vδ2^+^ γδ T cells (Vδ2-Jδ1/2/3/4), β2-microglobulin (β2M), and bacteria (PAN23S rDNA) (all from Integrated DNA Technologies, Coralville, IA) are shown in Supplementary Table 1. qPCR was performed using the KAPA Probe Fast qPCR kit (KAPA Biosystems, Wilmington, MA) and an ABI Prism ViiA7 (Applied Biosystems, Foster City, CA): 95°C for 3 min, then 40 cycles of 95°C for 3 sec and 60°C for 20 sec. Human DNA quantities were determined by amplifying the β2M gene [21]. The amount of Vα7.2-Jα12/20/33 and Vδ2-Jδ1/2/3/4 relative to β2M was determined by the comparative CT method (2^ΔΔ^CT). LinRegPCR was then used to calculate absolute β2M copy number [22] and this was multiplied by the relative amount of Vα7.2-Jα12/20/33 or Vδ2-Jδ1/2/3/4 to calculate the absolute abundance of MAIT or Vδ2^+^ γδ T cells (presented in arbitrary units). When no DNA was detected, a value less than the lowest amount detected was assigned (1 × 10^-10^ arbitrary units). To determine the bacterial load, 23S rDNA was detected using the KAPA SYBR FAST Universal kit (KAPA Biosystems): 95°C for 3 min, then 40 cycles of 95°C for 3 sec and 58°C for 30 sec. A standard curve with *S. aureus* DNA was used for quantification.

### Flow cytometry

PBMCs were stained with the following antibodies: CD3-PE/Cy7, Vδ2-FITC, Vα7.2-PE (BioLegend, San Diego, CA), CD8-eFluor450 (eBioscience, San Diego, CA), CD161-APC (Miltenyi Biotech, Bergisch Gladbach, Germany). Live/Dead Fixable Near IR dye (Life Technologies, Carlsbad, CA), and 123count eBeads (eBioscience) were included with each sample. Data was acquired on a FACSCanto II (BD Biosciences, San Jose, CA), and analysed using FlowJo version 10 software (Treestar, Inc., San Carlos, CA). The gating strategy is shown in Supplementary Figure 1. For comparisons of qPCR and flow cytometry, DNA was extracted from the same cryovial of PBMCs as analysed by flow cytometry. In some experiments MAIT cells and Vδ2^+^ T cells were sorted for DNA extraction using the BD FACSAria I (BD Biosciences).

### Analysis of Cytokine and Chemokine Production

Cytokine and chemokine levels in sputum samples were measured using the LEGENDplex 13-plex Human Inflammation Panel kit, measuring CCL2 (MCP-1), interferon-α (IFN-α), IFN-γ, interleukin-1β (IL-1β), IL-6, CXCL8 (IL-8), IL-10, IL-12 (p70), IL-17A, IL-18, IL-23, IL-33, and TNF-α, or the LEGENDplex 13-plex Human Proinflammatory Chemokine Panel Kit (BioLegend), measuring CCL2 (MCP-1), CCL3 (MIP-1α), CCL4 (MIP-1β), CCL5 (RANTES), CCL11 (exotoxin), CCL17 (TARC), CCL20 (MIP-3α), CXCL1 (GROα), CXCL5 (ENA-78), CXCL8 (IL-8), CXCL9 (MIG), CXCL10 (IP-10), and CXCL11 (I-TAC). Samples were acquired on a BD FACSCanto II, and analysed using LEGENDplex software (BioLegend).

### Metagenomics

16S rRNA sequencing was performed on sputum-extracted DNA at the Environmental Sample Preparation and Sequencing Laboratory at Argonne National Laboratories (Chicago, Illinois), as described in Supplementary Methods. Briefly, libraries of the V4 hypervariable region were prepared using primers 515F and 806R [23] and samples were sequenced on an Illumina Miseq using 2 × 250 bp read chemistry. The average number of reads per sample was 19,754 (range 1485-63,876). Quality control, assembly, and OTU assignment of sequences was done using Mothur [24], Taxonomy was assigned using the GreenGenes database version 13.5.99 [25]. Picrust was used to infer metagenomic capacity of 16S rRNA data [26]. Bacterial genes encoding for riboflavin and C5 isoprenoid biosynthesis were curated from the KEGG database (Supplementary Table 2).

### Statistics

Data were analysed in Prism 7 (GraphPad Software, San Diego, CA). Medians, interquartile range, and all data points are shown. Comparisons between 2 groups were made with the Mann-Whitney test. Comparisons between >2 groups were made with the Kruskal-Wallis test with Dunn’s multiple comparisons test. For continuous data, Spearman or Pearson correlations were calculated as indicated. Significance was defined as 2-sided P<0.05.

## Results

In this study, we sought to enumerate MAIT and Vδ2^+^ T cells at the site of infection in patients with CAP. As flow cytometric analysis was not possible due to the method of sample freezing, we used qPCR on DNA extracted from sputum samples to quantify innate-like lymphocyte (ILL)-specific VDJ TCR recombinations [21]. During VDJ recombination, variable and junctional segments are brought together in the genome to form a single exon. Here, we sought to develop a second generation qPCR assay capable of identifying the predominant MAIT cell TCRs (Vα7.2-Jα33/12/20) and a new assay to detect Vδ2^+^ TCRs (Vδ2-Jδ1/2/3/4) [27, 28].

To confirm the specificity of the primers and probes, qPCR was performed on DNA isolated from FACS-sorted PBMC populations. Significantly more Vα7.2-Jα12/20/33 and Vδ2-Jδ1/2/3/4 were detected in sorted MAIT cell and Vδ2^+^ T cell populations, respectively, than in depleted populations (Figure 1A-B). To determine the quantitative accuracy of the assays, we compared the absolute cell abundance of MAIT cells and Vδ2^+^ T cells in PBMCs as determined by flow cytometry and qPCR. Strong linear correlations for the numbers of MAIT and Vδ2^+^ T cells determined by qPCR and flow cytometry were observed (Figure 1C-D). Taken together, these results validate the qPCR method for measuring absolute cell abundance.

**Figure 1.**
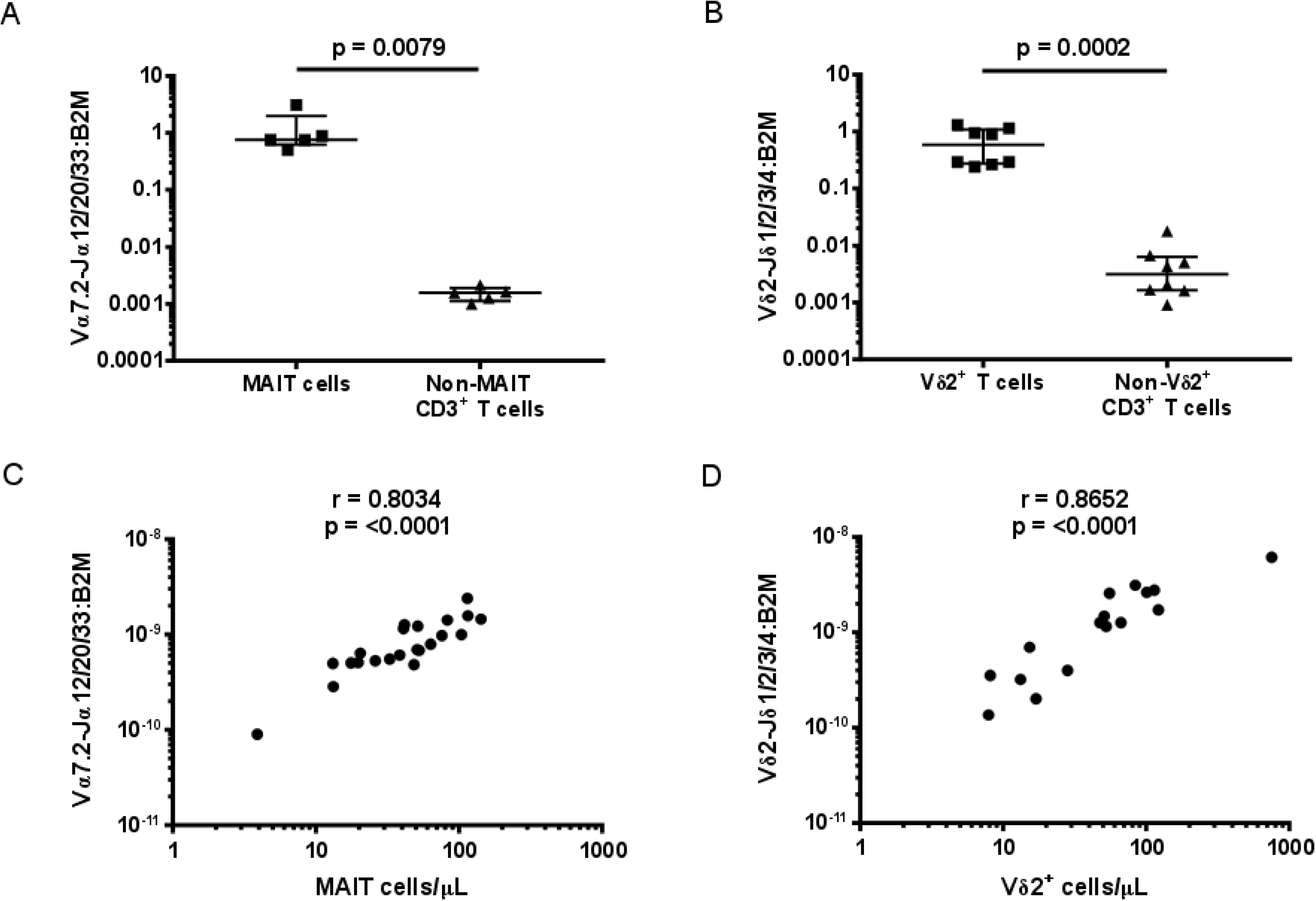
Validation of the qPCR method to determine MAIT and Vδ2^+^ T cell abundance. The specificity of the qPCR was determined by measuring the abundance of (A) Vα7.2-Jαl2/20/33 gDNA in MAIT cells (live/CD3^+^/CD8^+^/CD161^++^/Vα7.2^+^) and non-MAIT CD3^+^ T cells (live/CD3^+^/CD8^+^/CD161^-^/Vα7.2^-^), and (B) Vδ2-Jδl/2/3/4 gDNA in Vδ2^+^ cells (live/CD3^+^/Vδ2^+^) and Vδ2^-^ CD3^+^ T cells (Iive/CD3^+^/Vδ2^-^); each sorted from PBMCs and abundance expressed relative to β2M in arbitrary units. Correlation between the abundance of (C) MAIT and (D) Vδ2^+^ T cells measured by qPCR and by flow cytometry. (A) and (B) were analysed by Mann-Whitney tests. (C) and (D) were analysed using Spearman correlations. \*\**P*< 0.01, \*\*\**P* < 0.001 (Mann-Whitney test).

Sputum samples were available from 88 patients with CAP. Of these, 36 samples were excluded due to microscopic evidence of significant oropharyngeal contamination (> 10 squamous epithelial cells per 100x field). Of the 52 samples included in the study, 43 (83%) were expectorated and 9 were (17%) induced sputa. The median patient age was 67.5 (range 36 to 101) and 27 (52%) were female. Eighteen (35%) patients had pre-existing chronic obstructive pulmonary disease (COPD) or other structural lung disease and 11 (21%) had asthma. On chest x-ray, 31 (60%) had lobar consolidation while 21 (40%) had multilobar consolidation.

The abundance of MAIT cells, Vδ2^+^ T cells, and total innate-like lymphocytes (ILLs), defined as the sum of MAIT and Vδ2^+^ T cell abundance, were compared. MAIT and Vδ2^+^ T cell abundance in sputum were significantly correlated (Supplementary Figure 2A). There was no evidence of a correlation between MAIT cell, Vδ2^+^ T cell, or ILL abundance in sputum and patient age (Supplementary Figure 2B-D). MAIT cell, Vδ2^+^ T cell, and ILL abundance in sputum did not significantly differ by specimen type, gender, pre-existing lung disease, duration of illness prior to presentation, failure of outpatient antibiotic therapy, or type of consolidation on chest x-ray (Supplementary Table 3).

While there was no evidence of a difference in MAIT cell, Vδ2^+^ T cell, or total ILL abundance in patients requiring ICU admission (n=6) (Supplementary Figure 3), the abundance of MAIT cells and ILLs differed significantly with pneumonia severity as measured by the CURB65 score: patients with a CURB65 score of 0 had significantly more MAIT cells and total ILLs than those with a CURB65 score of 2 (Figure 2A, Supplementary Figure 4A). Patients with severe pneumonia by the IDSA/ATS criteria (n=4) had a lower MAIT cell abundance (Figure 2C), but total ILL abundance was not significantly different (Supplementary Figure 4B). There was no evidence of correlation between Vδ2^+^ T cell abundance and pneumonia severity, as calculated by the CURB65 or IDSA/ATS criteria (Figure 2B and 2D). Duration of stay in hospital was negatively correlated with ILL, MAIT cell, and Vδ2^+^ T cell abundance (Figure 3A-C).

**Figure 2.**
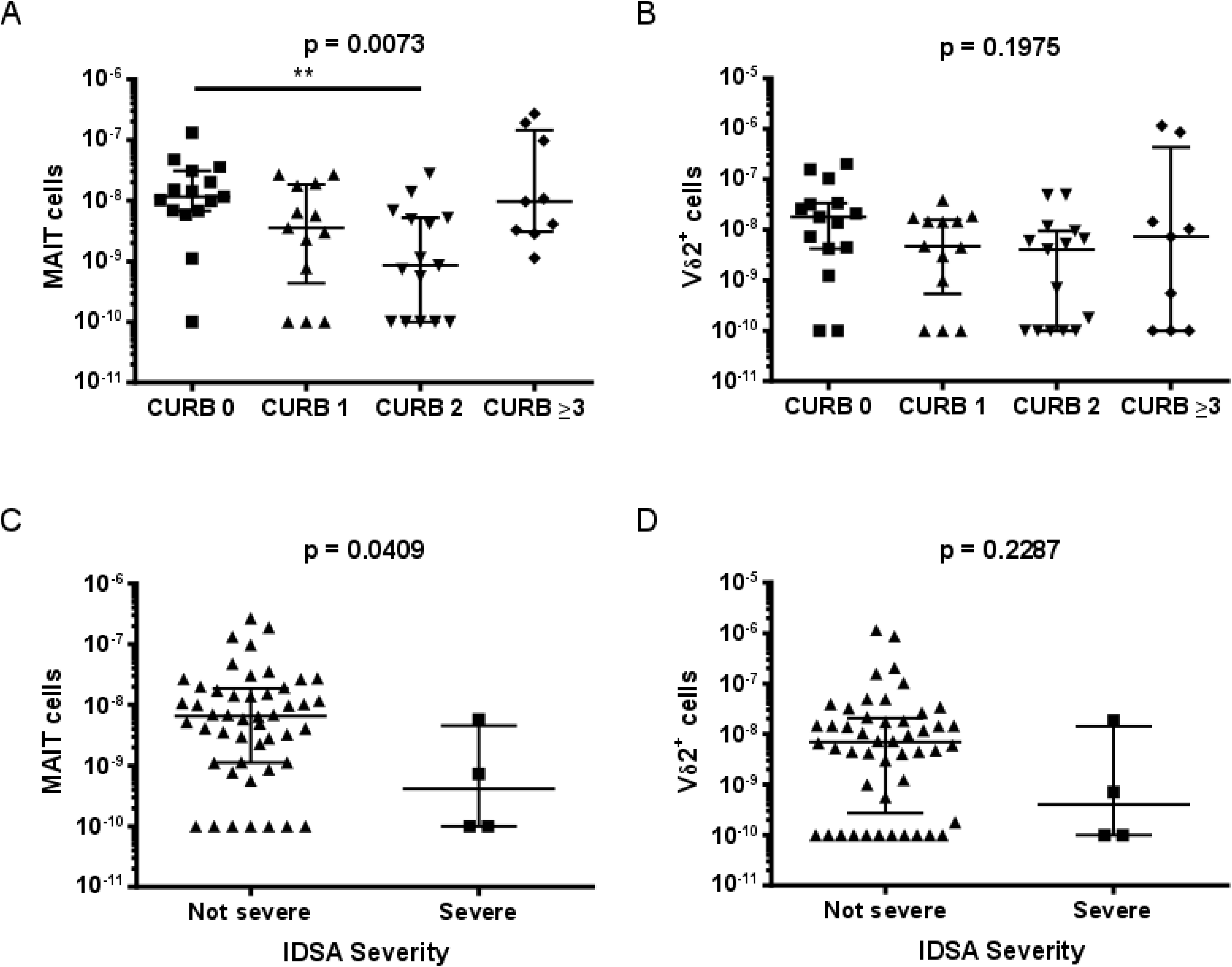
Reduced disease severity in patients with higher MAIT cell abundance in sputum. The abundance of (A) MAIT cells and (B) Vδ2^+^ T cells in sputum of patients by CURB-65 score. Abundance of (C) MAIT cells and (D) Vδ2^+^ T cells in sputum of patients by the IDSA severity criteria. MAIT cell and Vδ2^+^ T cell abundance were measured by qPCR and is expressed in arbitrary units. CURB-65 data was analysed using Kruskal-Wallis 1-way ANOVA and Dunn’s multiple comparison test; IDSA severity data was analysed by the Mann-Whitney test. \*\**P* < 0.01.

**Figure 3.**
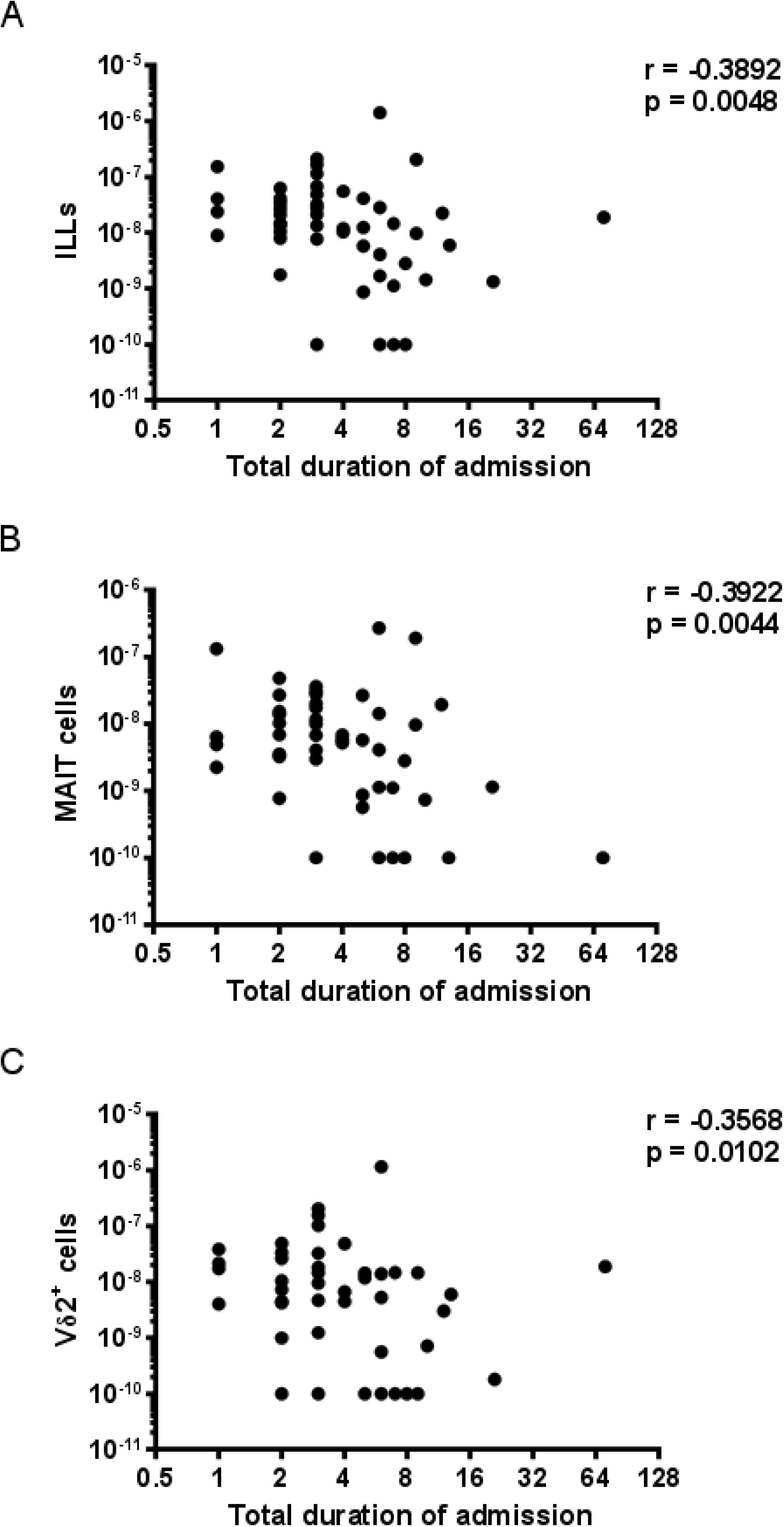
Duration of hospital admission is inversely correlated with the abundance of ILLs sputum. The abundance of (A) ILLs, (B) MAIT cells, and (C) Vδ2^+^ T cells were measured by qPCR (expressed in arbitrary units) and the association with duration of admission to hospital assessed with Spearman correlations.

MAIT cells, but not Vδ2^+^ T cells, were more abundant in sputum with more neutrophils, as determined by microscopy performed by the diagnostic laboratory prior to freezing (Supplementary Figure 5A-B). There was no evidence of a correlation between MAIT cell or Vδ2^+^ T cell abundance in sputum and the blood leukocyte count or C-reactive protein (CRP) levels (Supplementary Figure 5C-F).

A bacterial aetiology of infection was identified in 23 cases by routine diagnostic testing (Supplementary Table 4). MAIT cell, Vδ2^+^ T cell, and overall ILL abundance in sputum were all higher in patients with an identified bacterial pathogen (Figure 4A and 4B, Supplementary Figure 6A). Of note, no MAIT or Vδ2^+^ T cells were detected in several samples in which no bacterial pathogen was detected (Figure 4A and 4B). Vδ2^+^ T cell, but not MAIT cell, abundance was higher in patients with pneumonia caused by either *S. pneumoniae* or *Legionella* spp. (Supplementary Figure 6B-E). In contrast there was no indication of a correlation between either subset with total bacterial load (Figure 4C and 4D).

**Figure 4.**
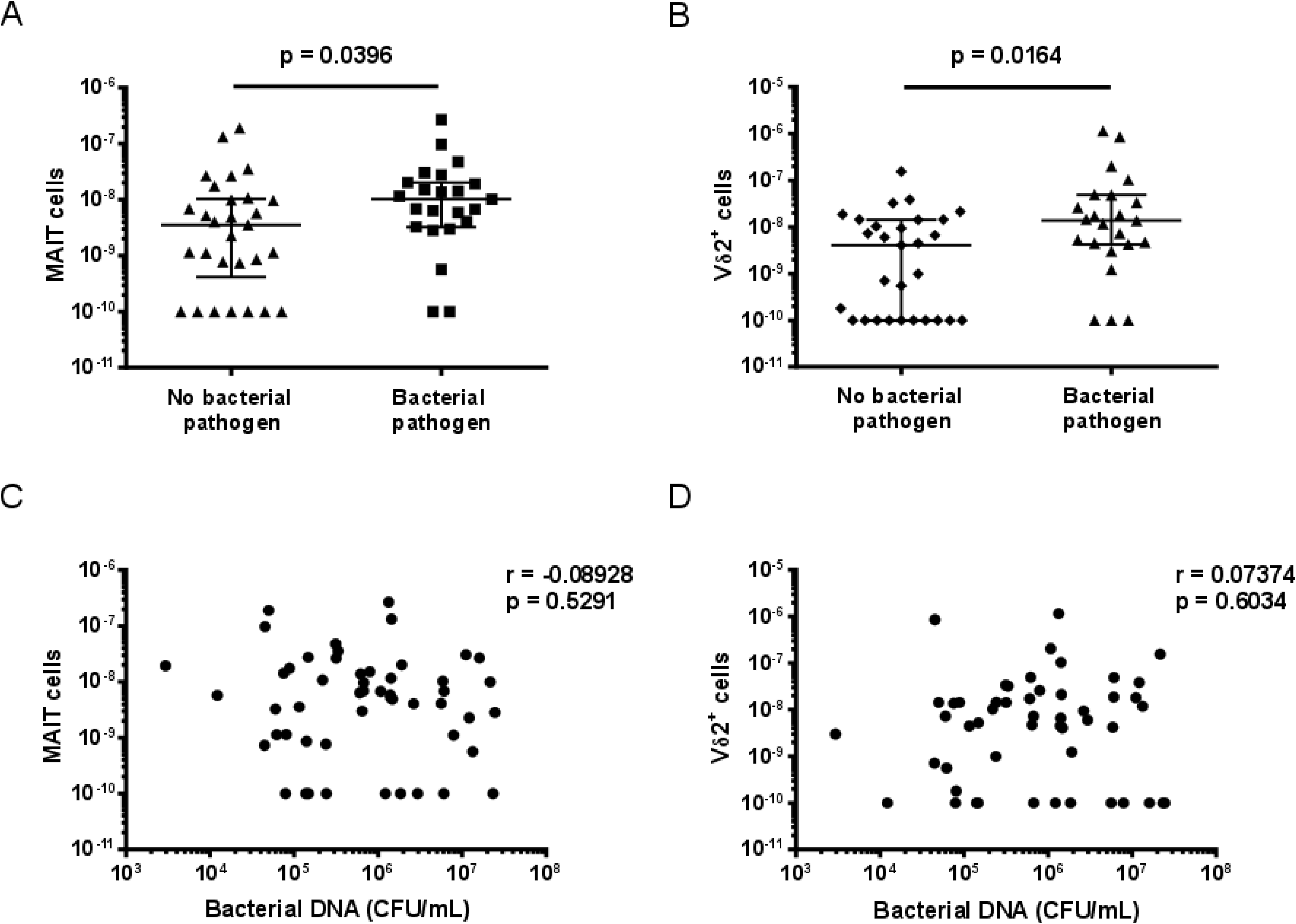
MAIT cell and Vδ2^+^ T cell abundance in sputum is higher in patients with an identified bacterial pathogen. The abundance of (A) MAIT cells and (B) Vδ2^+^ T cells in sputum of patients with and without an identified bacterial pathogen. The association between (C) MAIT cell and (D) Vδ2^+^ T cell abundance in sputum and total bacterial load. MAIT cell and Vδ2^+^ T cell abundance and total bacterial load were measured by qPCR and is expressed in arbitrary units. Bacterial pathogens were identified in the diagnostic laboratory by sputum culture, blood culture, PCR on sputum for *Legionella* spp., or by urinary antigen detection for *S. pneumoniae* or *Legionella pneumophila* serotype 1 (Supplementary Table 4). Differences between groups were analysed by Mann-Whitney tests (A, B). The association with total bacterial load was assessed with Spearman correlations (C, D). CFU/mL = colony forming units per millilitre.

As pneumonia can be a polymicrobial infection caused by components of the oral flora, we conducted 16S rRNA sequencing of samples to identify bacterial operational taxonomic units (OTUs) present in the sputum samples and predict the ability of those OTUs to make the metabolic ligands for ILLs. This revealed that there was no significant correlation in the inferred metagenomic data between the abundance of MAIT cells and the abundance of genes encoding for riboflavin synthesis, nor between the abundance of Vδ2^+^ T cells and the abundance of genes encoding for C5 isoprene synthesis (Supplementary Table 5). Culture of pathogens from sputum samples correlated with the abundance of genera containing pathogens in the 16S rRNA data (Supplementary Figure 7).

Finally, where there was sufficient sputum sample available, we analysed cytokine (n = 40) and chemokine (n=30) levels. The amount of IFN-γ in sputum was significantly correlated with MAIT cell, Vδ2^+^ T cell, and overall ILL abundance (Table 1, Supplementary figure 8A-C). The amount of IFN-α in sputum was significantly correlated with MAIT cell abundance alone (Table 1, Supplementary Figure 8D-F). The amount of CXCL11 in sputum was significantly correlated with Vδ2^+^ T cell and overall ILL abundance but not with MAIT cell abundance (Table 1, Supplementary figure 8G-I). No other cytokines or chemokines analysed displayed evidence of a significant correlation with MAIT cell, Vδ2^+^ T cell, or overall ILL abundance (Table 1).

**Table 1.**

| Analyte | ILL |  |  | MAIT cells |  |  | Vδ2+ T cells |  |  |
| --- | --- | --- | --- | --- | --- | --- | --- | --- | --- |
|  | Spearman r | 95% CI | P (two-tailed) | Spearman r | 95% CI | P (two-tailed) | Spearman r | 95% CI | P (two-tailed) |
| CCL2* | 0.026 | -0.347 to 0.392 | 0.891 | -0.031 | -0.397 to 0.343 | 0.869 | 0.098 | -0.283 to 0.451 | 0.608 |
| CCL2** | 0.167 | -0.162 to 0.463 | 0.303 | 0.171 | -0.158 to 0.465 | 0.293 | 0.201 | -0.128 to 0.489 | 0.215 |
| CCL3* | 0.030 | -0.344 to 0.396 | 0.875 | 0.106 | -0.275 to 0.458 | 0.577 | -0.029 | -0.395 to 0.345 | 0.879 |
| CCL4* | -0.038 | -0.402 to 0.337 | 0.842 | 0.071 | -0.307 to 0.430 | 0.710 | -0.101 | -0.454 to 0.280 | 0.596 |
| CCL5* | 0.220 | -0.164 to 0.545 | 0.243 | 0.251 | -0.131 to 0.568 | 0.181 | 0.088 | -0.292 to 0.444 | 0.644 |
| CCL11* | 0.027 | -0.346 to 0.393 | 0.888 | 0.120 | -0.261 to 0.469 | 0.527 | -0.116 | -0.466 to 0.265 | 0.540 |
| CCL17* | -0.192 | -0.525 to 0.191 | 0.309 | -0.243 | -0.562 to 0.140 | 0.196 | -0.248 | -0.566 to 0.134 | 0.186 |
| CCL20* | 0.153 | -0.230 to 0.495 | 0.419 | 0.168 | -0.216 to 0.506 | 0.376 | 0.127 | -0.255 to 0.474 | 0.505 |
| CXCL1* | 0.018 | -0.355 to 0.385 | 0.927 | 0.077 | -0.302 to 0.435 | 0.686 | 0.006 | -0.365 to 0.375 | 0.975 |
| CXCL5* | -0.169 | -0.507 to 0.214 | 0.372 | -0.197 | -0.528 to 0.187 | 0.297 | -0.118 | -0.468 to 0.263 | 0.534 |
| CXCL8* | -0.053 | -0.415 to 0.323 | 0.780 | 0.031 | -0.343 to 0.396 | 0.872 | -0.016 | -0.383 to 0.356 | 0.935 |
| CXCL8** | -0.040 | -0.356 to 0.284 | 0.806 | 0.073 | -0.253 to 0.384 | 0.653 | 0.006 | -0.314 to 0.326 | 0.969 |
| CXCL9* | 0.051 | -0.325 to 0.413 | 0.790 | -0.074 | -0.432 to 0.304 | 0.697 | 0.095 | -0.285 to 0.449 | 0.618 |
| CXCL10* | 0.154 | -0.229 to 0.496 | 0.417 | -0.112 | -0.463 to 0.269 | 0.554 | 0.240 | -0.143 to 0.560 | 0.202 |
| CXCL11* | **0.376** | **0.007 to 0.655** | **0.041** | 0.165 | -0.218 to 0.504 | 0.383 | **0.392** | **0.025 to 0.665** | **0.032** |
| IFNα** | 0.311 | -0.010 to 0.574 | 0.051 | **0.328** | **0.009 to 0.587** | **0.039** | 0.230 | -0.097 to 0.513 | 0.153 |
| IFNγ** | **0.488** | **0.200 to 0.699** | **0.001** | **0.421** | **0.117 to 0.653** | **0.007** | **0.487** | **0.198 to 0.698** | **0.001** |
| IL1β** | 0.200 | -0.129 to 0.489 | 0.217 | 0.296 | -0.026 to 0.563 | 0.063 | 0.083 | -0.243 to 0.393 | 0.609 |
| IL6** | 0.123 | -0.205 to 0.427 | 0.449 | 0.086 | -0.241 to 0.395 | 0.599 | 0.183 | -0.146 to 0.475 | 0.258 |
| IL10** | 0.067 | -0.259 to 0.379 | 0.684 | 0.176 | -0.153 to 0.469 | 0.278 | 0.079 | -0.248 to 0.389 | 0.630 |
| IL12p70** | 0.171 | -0.158 to 0.466 | 0.291 | 0.244 | -0.083 to 0.523 | 0.130 | 0.062 | -0.264 to 0.374 | 0.705 |
| IL17A** | 0.251 | -0.076 to 0.528 | 0.119 | 0.256 | -0.070 to 0.532 | 0.111 | 0.239 | -0.088 to 0.519 | 0.138 |
| IL18** | 0.214 | -0.114 to 0.500 | 0.186 | 0.197 | -0.132 to 0.486 | 0.224 | 0.241 | -0.086 to 0.521 | 0.134 |
| IL23** | -0.012 | -0.331 to 0.310 | 0.943 | 0.103 | -0.224 to 0.410 | 0.526 | -0.042 | -0.357 to 0.282 | 0.796 |
| IL33** | 0.200 | -0.128 to 0.490 | 0.216 | 0.164 | -0.164 to 0.460 | 0.311 | 0.234 | -0.093 to 0.515 | 0.146 |
| TNFα** | 0.282 | -0.042 to 0.552 | 0.078 | 0.287 | -0.036 to 0.556 | 0.072 | 0.249 | -0.077 to 0.527 | 0.121 |

IFN, interferon; IL, interleukin; TNF, tumour necrosis factor. Significant P values are bolded.
\* Measured with LEGENDplex 13-plex Human Proinflammatory Chemokine Panel kit (BioLegend) (n=30)
\*\* Measured with LEGENDplex 13-plex Human Inflammation Panel kit (BioLegend) (n=40)

## Discussion

In this study, we quantified MAIT cells and Vδ2^+^ T cells at the site of infection in patients with CAP and found their abundance to be inversely associated with clinical markers of severity. Higher numbers of MAIT cells and Vδ2^+^ T cells were found in sputum of CAP patients with a confirmed bacterial aetiology, but their abundance was not related to bacterial load or the predicted capacity of bacteria to produce the MAIT cell or Vδ2^+^ T cell activating ligands. Instead, Vδ2^+^ T cell abundance correlated with the concentration of CXCL11, MAIT cell abundance with the concentration of IFN-α, and both cell populations with the amount of IFN-γ in sputum. Overall, this suggests an important role for MAIT cells and Vδ2^+^ T cells in the immune response to CAP.

As the frozen sputum specimens were not suitable for flow cytometry, we developed a molecular assay to quantify MAIT and Vδ2^+^ γδ T cell DNA. By using multiple junctional primers, this method can detect the most common recombined MAIT cell TCRs and all possible Vδ2^+^ γδ T cell TCRs [27, 28]. Importantly, the abundance of MAIT and Vδ2^+^ cells determined by qPCR and flow cytometry were strongly correlated. This method could be used to complement flow cytometry and enable quantification of ILLs in a range of tissues and body fluids.

The abundance of MAIT and Vδ2^+^ T cells in sputum of patients with CAP was highly correlated. This suggests that common signals lead to the recruitment and/or proliferation of these ILLs. Indeed, both cell populations express chemokine receptors associated with trafficking to the lungs and sites of inflammation [3, 17, 29], and have the ability to proliferate [17, 30].

While the levels of ILLs were higher in those with an identified bacterial pathogen, there was no evidence of association with total bacterial load. An association between MAIT cell recruitment to the lungs and bacterial inoculum has recently been reported in a mouse model of *Legionella longbeachae* pneumonia [10]. The lack of correlation between total bacterial load and MAIT cell, Vδ2^+^ T cell, or ILL abundance in our study may reflect heterogeneity in the duration of infection, prior antibiotic treatment, sample contamination with oropharyngeal microflora, and non-bacterial causes of pneumonia. Indeed, the most commonly detected pathogens in pneumonia are viruses [31–34], As this study was conducted outside the influenza season [35], patients were not tested for respiratory viruses; however, CAP resulting from other respiratory viruses, such as human rhinovirus, cannot be excluded [31]. Identification of the causative agent in CAP is notoriously difficult; in a recent pneumonia aetiology study, extensive testing identified a pathogen in only 38% of patients [34]

Interestingly, not all bacterial pathogens identified are able to produce the ligands for MAIT and Vδ2^+^ T cells. Vδ2^+^ T cell abundance was higher in those with *S. pneumoniae* or *Legionella* infection, yet these species do not produce HMB-PP [36]. Further, neither MAIT cell nor Vδ2^+^ T cell abundance correlated with the quantity of bacteria in sputum with the capacity to produce riboflavin or HMB-PP respectively (as inferred from metagenomic analysis). Therefore, the abundance of TCR ligands is unlikely to be the sole determinant of MAIT and Vδ2^+^ T cell recruitment and/or proliferation, although their concentrations have not been measured. The abundance of Vδ2^+^ T cells in sputum correlated with the concentration of CXCL11 (I-TAC), which is produced by monocytes, endothelial cells, and fibroblasts in response to IFN-γ and IFN-β, and binds to CXCR3 [37], CXCR3 is expressed by Vδ2^+^ T cells and their chemotaxis in response to CXCR3 ligands has previously been reported [38]. While the correlation of Vδ2^+^ T cell abundance with CXCL11 concentration may be confounded by the concentration of IFN-γ, the lack of correlation of MAIT cell abundance with CXCL11 concentration argues against this. While MAIT cells express CXCR3, they also express high levels of CD26, which has been shown to inactivate CXCL11 and prevent T cell chemotaxis [39, 40]. No chemokines were identified that correlated with MAIT cell abundance. However, IFN-α concentration correlated with MAIT cell abundance but not Vδ2^+^ T cell abundance. IFN-α2 has been shown to be a chemotactic factor for T cells [41]. Alternatively, IFN-α may result in activation and proliferation of MAIT cells in the lung [42, 43]. In a mouse model of influenza infection, accumulation of MAIT cells in the lungs was dependent upon IL-18 [11]. However, in our study, MAIT cell abundance was not associated with IL-18 concentration.

MAIT cell abundance was positively correlated with neutrophil infiltration. Neutrophils are essential in the control of multiple bacterial pulmonary infections, and have a major role in the initiation of the adaptive immune response [44], MAIT cells produce IL-17A, inducing production of CXCL-8, resulting in neutrophil recruitment [45, 46]. Although IL-17A was detected in the sputum samples, the amount did not significantly correlate with MAIT cell abundance. Similarly, while γδ T cells have been found to produce IL-17 in response to lung infection [47], we did not detect a significant correlation between the abundance of Vδ2^+^ T cells and IL-17A production or neutrophil abundance. This suggests that other cell types may contribute to the production of IL-17A and CXCL-8, and hence to neutrophil recruitment. The correlation between MAIT cell and neutrophil abundance may reflect co-recruitment in response to inflammation, with the role of MAIT cells in recruiting neutrophils uncertain.

We found that the levels of IFN-γ in sputum were positively correlated with MAIT cell, Vδ2^+^ T cell, and overall ILL abundance in sputum, suggesting these cells are a major source of IFN-γ in CAP. Inflammatory cytokines, such as IFN-γ, TNF-α, and IL-17, are critical effector cytokines in pulmonary infection. In particular, IFN-γ has an important role in the response to various pulmonary bacterial infections [48, 49]. It has recently been reported that immune defence in pulmonary infection with *L. longbeachae* in Rag2^-/-^γC^-/-^ mice is reliant upon IFN-γ production by adoptively transferred MAIT cells; a significant reduction in survival was shown in mice where adoptively transferred MAIT cells were from IFN-γ^-/-^ mice [10]. However, mortality was unchanged when MAIT cells from TNF^-/-^, IL-17^-/-^, perform^-/-^ or granzyme A and B^-/-^ mice were transferred [10]. Production of IFN-γ, TNF-α, and IL-17 by MAIT and γδ T cells has previously been reported in mouse lung infection models [50, 51].

The abundance of MAIT cells and Vδ2^+^ cells in sputum was inversely associated with the severity of infection. Higher numbers of MAIT cells were found in the sputum of patients with low severity pneumonia (CURB65 score and IDSA/ATS criteria). CURB65 is a clinical scoring system that allows the stratification of patients into mortality risk groups [52], The IDSA/ATS criteria are designed to identify patients who require management in an intensive care or high dependency unit [53]. While MAIT cell abundance was significantly lower in patients with a CURB65 score of 2 than in patients with a score of 0, the difference between a CURB65 score of ≥3 and 0 was not significant. This may be due to the low numbers of patients with severe pneumonia in this cohort or greater recruitment of MAIT cells to the lungs in patients with the most severe infections. There was a significant difference in MAIT cell abundance between those with non-severe and severe pneumonia, as classified by the IDSA/ATS criteria, however no difference was seen with ICU admission. Again, this may be due to low numbers of severe cases or other reasons for subsequent ICU admission. Both MAIT cell and Vδ2^+^ T cell numbers in sputum were negatively correlated with the duration of hospital admission. Duration of admission may reflect severity of infection but could be confounded by comorbidities, such as COPD and age [54], MAIT cells numbers in the blood decrease with age [55] and are also depleted from the blood and lungs of patients with COPD [56]. However, neither MAIT cell nor Vδ2^+^ T cell abundance in sputum was correlated with age and there was no difference between those with or without COPD (data not shown). In support of these findings, MAIT cells have recently been shown to have a non-redundant role in protecting against pulmonary infection with *L. longbeachae*; bacterial clearance was diminished in MAIT cell deficient mice and enhanced in mice with an expanded MAIT cell population [10].

Overall, our findings suggest that ILLs, particularly MAIT cells, play a protective role against severe infection. This is consistent with the previous finding that MAIT cell numbers are reduced in the blood of severely unwell patients admitted to intensive care, especially those with a bacterial infection, and that MAIT cell recovery may protect against subsequent nosocomial infection [13]. In contrast, it has been suggested that MAIT and Vδ2^+^ T cells contribute to a poorer clinical outcome in patients with a first episode of continuous ambulatory peritoneal dialysis (CAPD) peritonitis [57], Of note, other bacterial factors, such as endotoxins, virulence factors, ability to form biofilms, and resistance to antimicrobials may influence inflammatory cytokine production and the failure rate of peritoneal dialysis and were not assessed in that study. Therefore, ILLs and the inflammatory response they induce may be protective in one clinical scenario (pneumonia) but deleterious in another (CAPD peritonitis).

Several limitations should be noted. Firstly, the method of sample preservation excluded the possibility of immunophenotyping. Secondly, MAIT cells and Vδ2^+^ T cells were not detected in several samples, falling below the limit of detection of the assay; this prevented us from fitting multiple regression models. Thirdly, no paired blood samples were available to compare ILL abundance. Thirdly, as discussed above, this cohort lacked severe cases. Fourthly, the abundance of ILLs in the airways of healthy controls or patients with stable asthma or COPD was not assessed; Vα7.2-Jα33 mRNA can be detected in in induced sputum samples from healthy volunteers, and at higher levels than in patients with stable COPD or an acute exacerbation of COPD [58]. Fifthly, we have not made an adjustment for multiple testing in our analysis, so replication in a prospective cohort will be important to assess the hypotheses generated by this study. Finally, there may be unmeasured confounders that account for our findings; therefore it is not possible to determine causation.

In conclusion, we have developed a method to quantify MAIT and Vδ2^+^ T cell semi-invariant TCR DNA. Using this method, we confirmed the presence of ILLs at the site of infection, and have identified an association of these cells with an improved clinical outcome. This should be investigated further in prospective studies. If these findings are confirmed, immunotherapies to enhance MAIT cell numbers and function could be considered for the prevention of severe CAP.

## Supporting information

## Acknowledgements

This work was supported by a University of Otago Research Grant.

JEU, DRM and XCM conceived the study; JEU, RFH, DRM, and XCM designed the study protocol; RFH conducted the experiments; RFH, XW, XCM, MS, and JEU carried out the analysis and interpretation of data; MRS provided statistical advice; RFH and MS drafted the manuscript; RFH, MS, and JEU critically revised the manuscript for intellectual content. All authors read and approved the final manuscript.

## Conflicts of Interest

The authors have no conflicts of interest.

---

**Supplementary Figure 1. Gating strategy to determine MAIT and Vδ2^+^ T cell counts for comparison to qPCR data.**

**Supplementary Figure 2. MAIT cell and Vδ2^+^ T cell abundance in sputum are correlated, but no correlation with age is seen.** MAIT cell and Vδ2^+^ T cell abundance in sputum of patients with CAP was measured by qPCR and is expressed in arbitrary units. (A) The relationship between MAIT cell and Vδ2^+^ T cell abundance. (B-D) The association between (B) ILL, (C) MAIT cell, and (D) Vδ2^+^ T cell abundance in sputum and patient age. Relationships between variables were assessed with Spearman correlations.

**Supplementary Figure 3. Abundance of ILLs does not differ with ICU admission.** Abundance of (A) ILLs, (B) MAIT cells, and (C) Vδ2^+^ T cells in sputum of patients with CAP requiring admission to the intensive care unit (ICU) or not. Cell abundance is expressed in arbitrary units. Data were analysed by Mann-Whitney tests.

**Supplementary Figure 4. Reduced disease severity in patients with higher ILL abundance in sputum.** (A) The abundance of ILLs in sputum of patients by CURB-65 score. (B) The abundance of ILLs in severe cases of pneumonia, as defined by the IDSA severity criteria. Cell abundance is expressed in arbitrary units. CURB-65 data was analysed using Kruskal-Wallis 1-way ANOVA and Dunn’s multiple comparison test; IDSA severity data was analysed by the Mann-Whitney test. \**P* < 0.05.

**Supplementary Figure 5. MAIT cell abundance in sputum is associated with the number of neutrophils in the sputum, but not with blood leukocyte count or with CRP.** The abundance of (A) MAIT cells and (B) Vδ2^+^ T cells by the number of neutrophils per low powered field (LPF) in sputum. The association of (C) MAIT cell and (D) Vδ2^+^ T cell abundance in sputum with blood leukocyte count, The association of (E) MAIT cell and (F) Vδ2^+^ T cell abundance in sputum with blood CRP concentration (mg/L). MAIT cell and Vδ2^+^ T cell abundance was determined by qPCR and is expressed in arbitrary units. Microscopy was performed on sputum in the diagnostic laboratory prior to freezing. Neutrophil data was analysed with the Kruskal-Wallis 1-way ANOVA. Blood leukocyte and CRP data were analysed using Spearman correlations. WCC = white cell (leukocyte) count; ns = not significant; CRP = C-reactive protein.

**Supplementary Figure 6. Vδ2^+^ T cell abundance in sputum is higher in patients with CAP caused by *Streptococcus pneumoniae* **or** *Legionella* spp.** (A) The abundance of ILLs in sputum of patients with and without an identified bacterial pathogen. (B-C) The abundance of MAIT cells in sputum of patients with CAP caused by (B) *S. pneumoniae* or (C) *Legionella* spp. The abundance of Vδ2^+^ T cells in sputum of patients with CAP caused by (D) *S. pneumoniae* or (E) *Legionella* spp. Cell abundance is expressed in arbitrary units. Differences between groups were analysed by Mann-Whitney tests.

**Supplementary Figure 7. Relationship between cultured pathogens and 16S rRNA data.** Genera of clinical interest are indicated on the X axis, while the Y axis shows the log relative abundance of each genera of interest within the 16S rRNA data. Facets correspond to groups of samples from which pathogen was cultured (HFLU = *Haemophilus influenzae* (n=5); LGN = *Legionella* spp. (n=14); MCAT = *Moraxella catarrhalis* (n=5); PAER = *Pseudomonas aeruginosa* (n=1); PNEU = *Streptococcus pneumoniae* (n=4); SAUR = *Staphylococcus aureus* (n=1); NSG = no significant growth (n=41)). The mean abundance of genera is not significantly different between groups, as measured by the Kruskal-Wallis and Wilcoxon signed-rank tests, except for between samples +/- *Legionella* (p<0.001).

**Supplementary Figure 8. Association of ILL abundance in sputum with sputum concentrations of IFN-γ, IFN-α, and CXCL11.** (A-C) The association of sputum IFN-γ concentrations with the abundance of (A) ILLs, (B) MAIT cells, and (C) Vδ2^+^ T cells in sputum. (D-F) The association of sputum IFN-α concentrations with the abundance of (D) ILLs, (E) MAIT cells, and (F) Vδ2 ^+^ T cells in sputum. (G-I) The association of sputum CXCL11 concentrations with the abundance of (G) ILLs, (H) MAIT cells, and (I) Vδ2^+^ T cells in sputum. Cell abundance was determined by qPCR and is expressed in arbitrary units. IFN-γ, IFN-α, and CXCL11 were measured by LEGENDplex bead array. Associations between different cell populations and soluble mediators were assessed with Spearman correlations.

## Supplementary Methods

### Protocol for 16S rRNA amplicon library preparation and sequencing

Briefly, PCR amplicon libraries targeting the 16S rRNA encoding gene present in metagenomic DNA are produced using a barcoded primer set adapted for the Illumina HiSeq2500 and MiSeq [23]. DNA sequence data is then generated using Illumina paired-end sequencing at the Environmental Sample Preparation and Sequencing Facility (ESPSF) at Argonne National Laboratory. Specifically, the V4 region of the 16S rRNA gene (515F-806R) is PCR amplified with region-specific primers that include sequencer adapter sequences used in the Illumina flow cell [23, 61]. The forward amplification primer also contains a twelve base barcode sequence that supports pooling of up to 2,167 different samples in each lane [23, 61]. Each 25 μL PCR reaction contains 9.5 μL of MO BIO PCR Water (Certified DNA-Free), 12.5 μL of QuantaBio’s AccuStart II PCR ToughMix (2x concentration, 1x final), 1 μL Golay barcode tagged Forward Primer (5 μM concentration, 200 pM final), 1 μL Reverse Primer (5 μM concentration, 200 pM final), and 1 μL of template DNA. The conditions for PCR are as follows: 94°C for 3 minutes to denature the DNA, with 35 cycles at 94°C for 45 s, 50°C for 60 s, and 72°C for 90 s; with a final extension of 10 min at 72°C to ensure complete amplification. Amplicons are then quantified using PicoGreen (Invitrogen) and a plate reader (Infinite^®^ 200 PRO, Tecan). Once quantified, volumes of each of the products are pooled into a single tube so that each amplicon is represented in equimolar amounts. This pool is then cleaned up using AMPure XP Beads (Beckman Coulter), and then quantified using a fluorometer (Qubit, Invitrogen). After quantification, the molarity of the pool is determined and diluted down to 2 nM, denatured, and then diluted to a final concentration of 6.75 pM with a 10% PhiX spike for sequencing on the Illumina MiSeq. Amplicons are sequenced on a 251bp x 12bp x 251bp MiSeq run using customized sequencing primers and procedures [23].

### Taxonomic Assignment

Data was processed according to the authors’ recommended best practices at: https://www.mothur.org/wiki/MiSeq_SOP. Assembled sequences longer than 275 bp or with ambiguities were removed during quality filtering. For taxonomic assignment, sequences were aligned to the greengenes reference database (version gg_13_5_99) [25], using a 97% identity threshold.

