## Supplementary figures and images for "Role of Innate-like Lymphocytes in the Pathogenesis of Community Acquired Pneumonia"

### Supplementary file 1

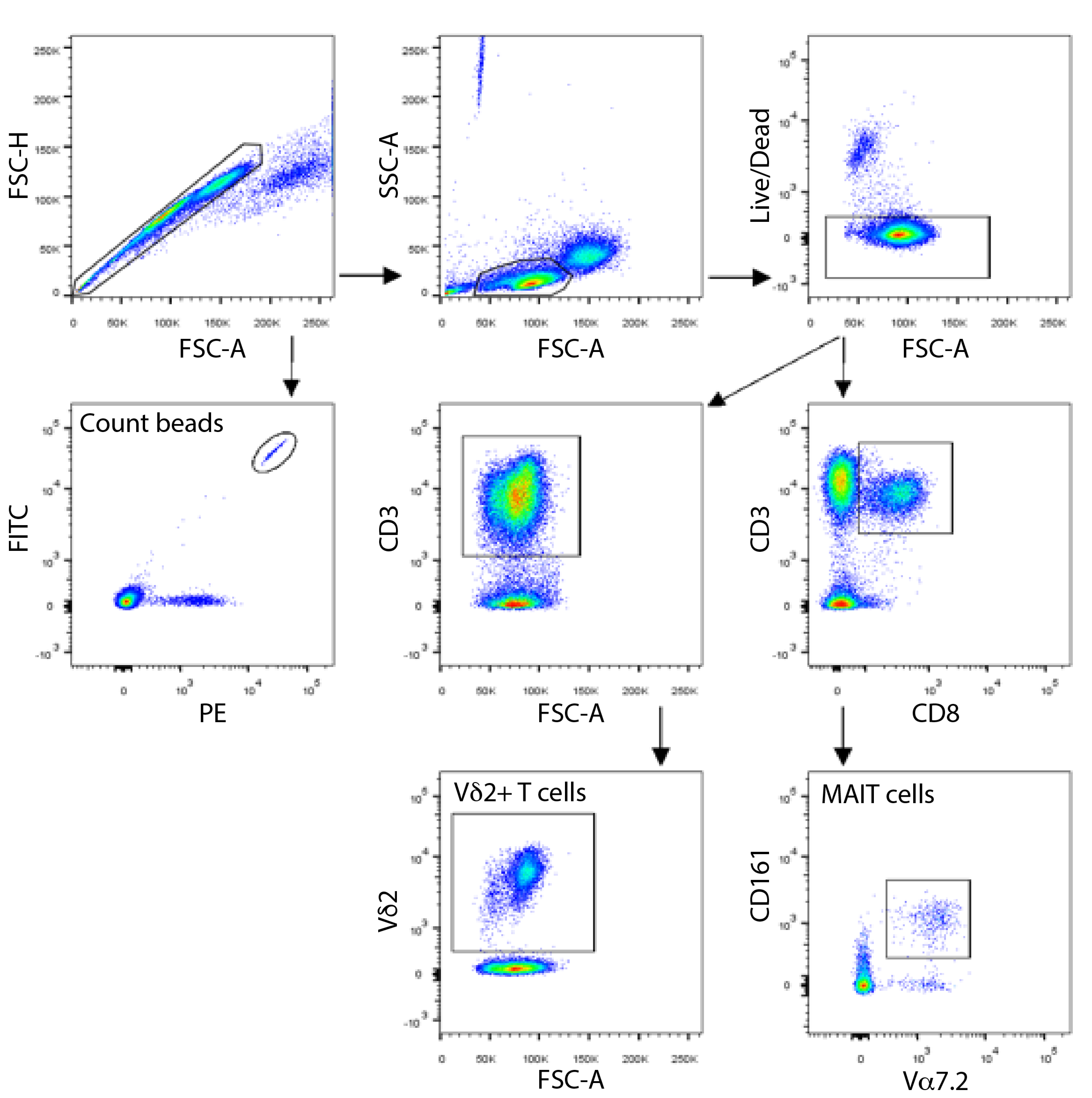

### Supplementary file 2

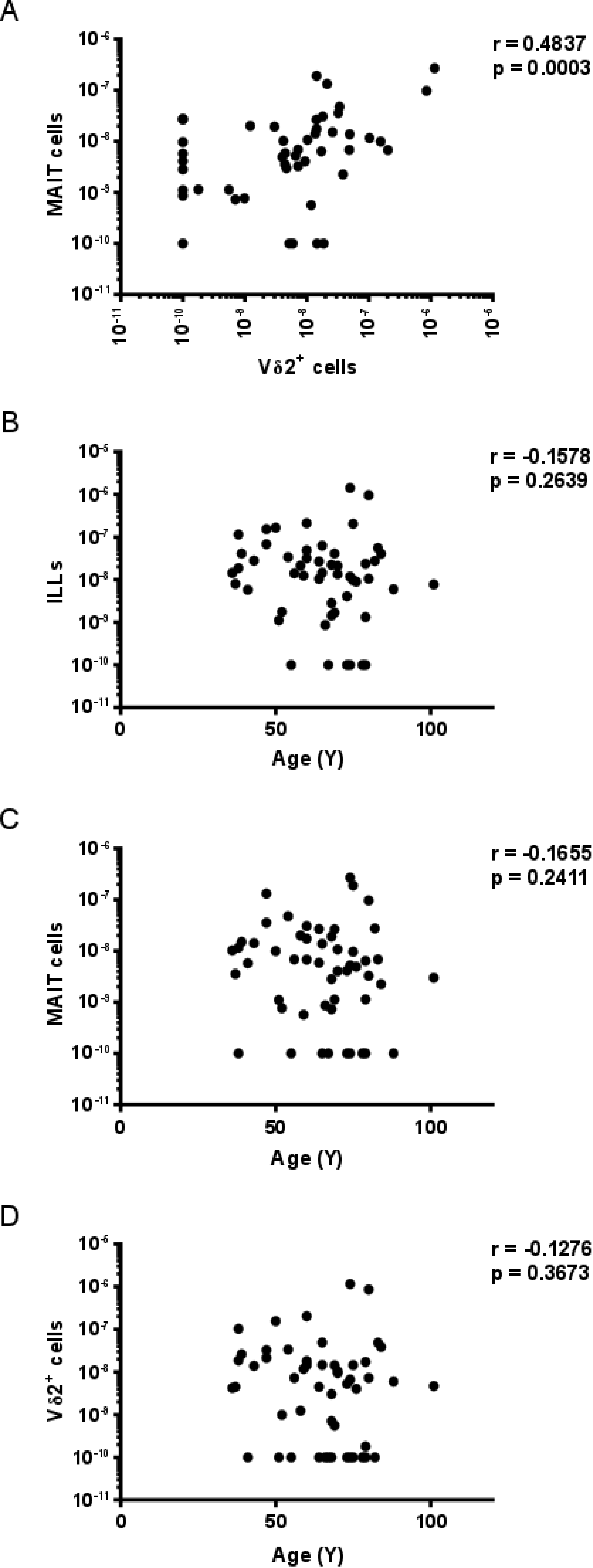

### Supplementary file 3

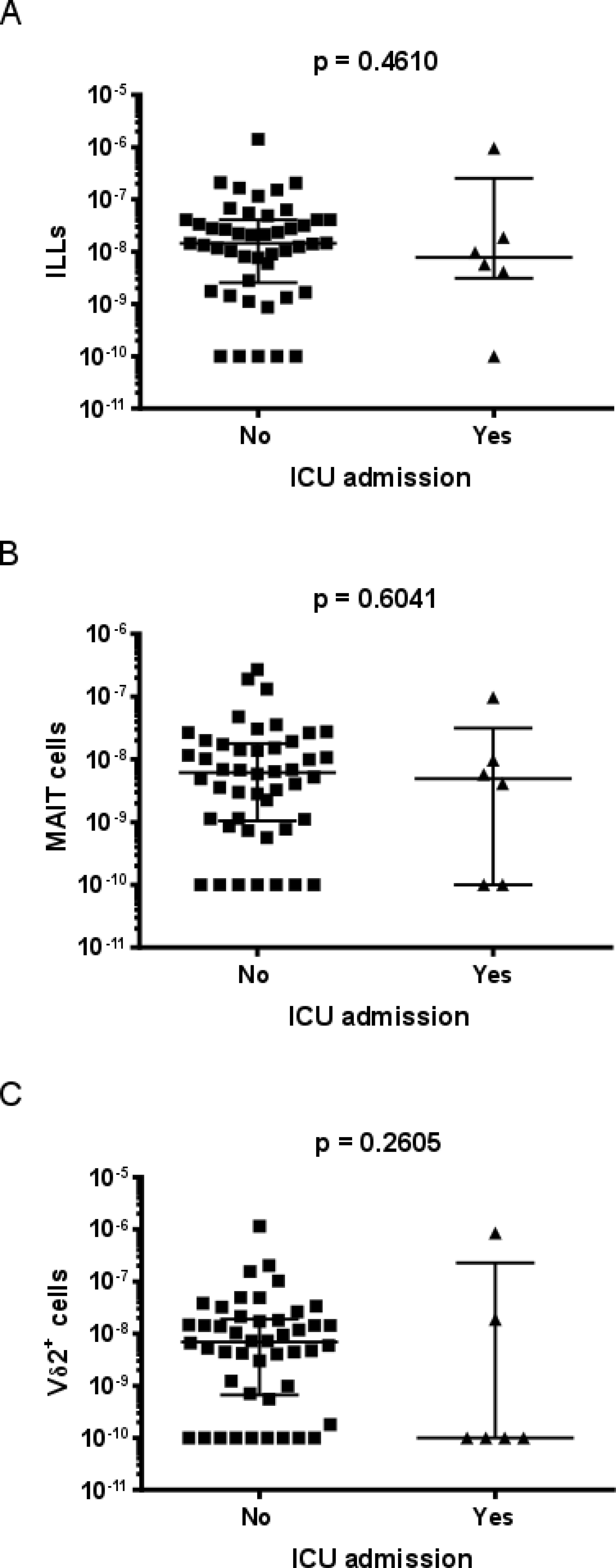

### Supplementary file 4

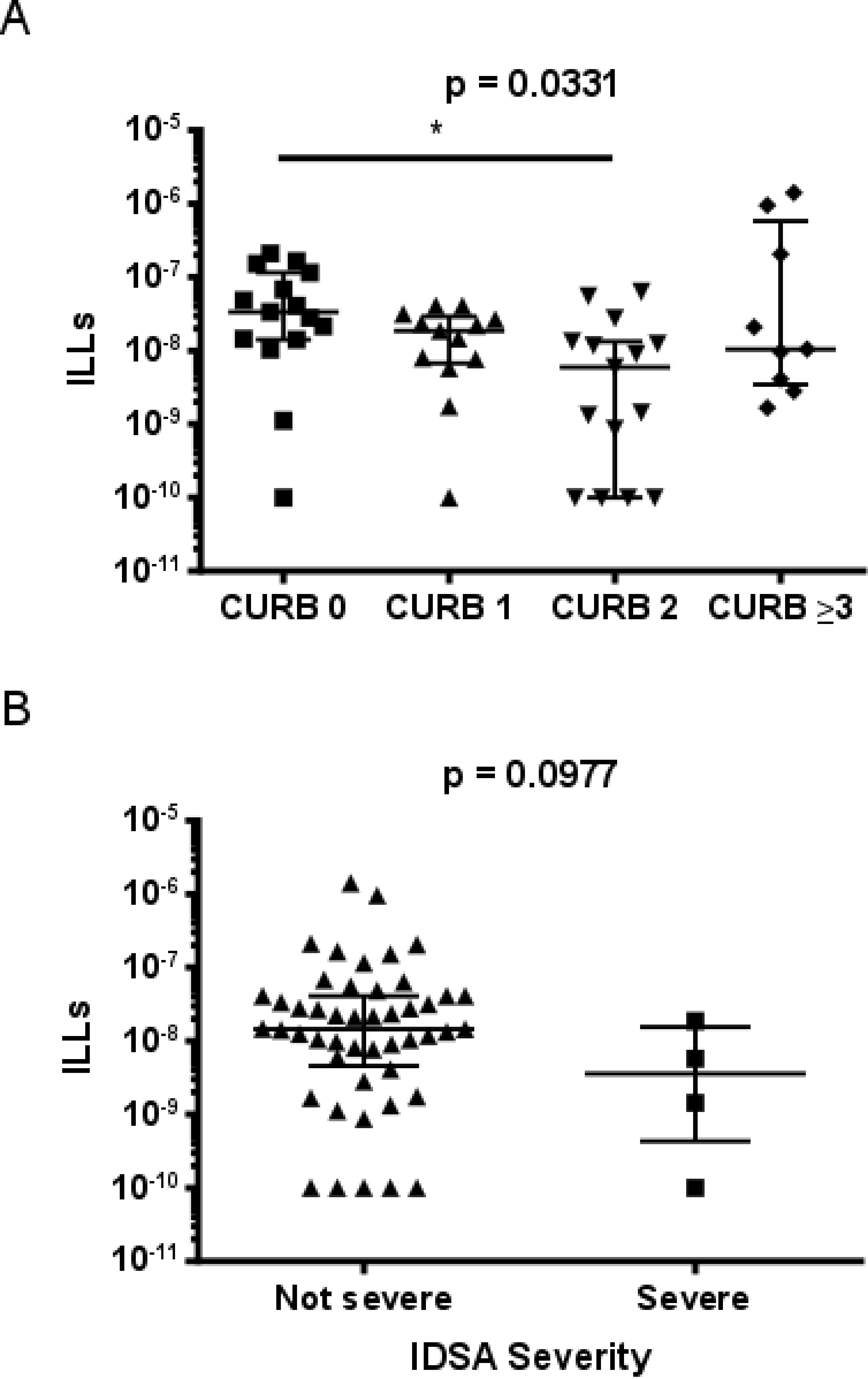

### Supplementary file 5

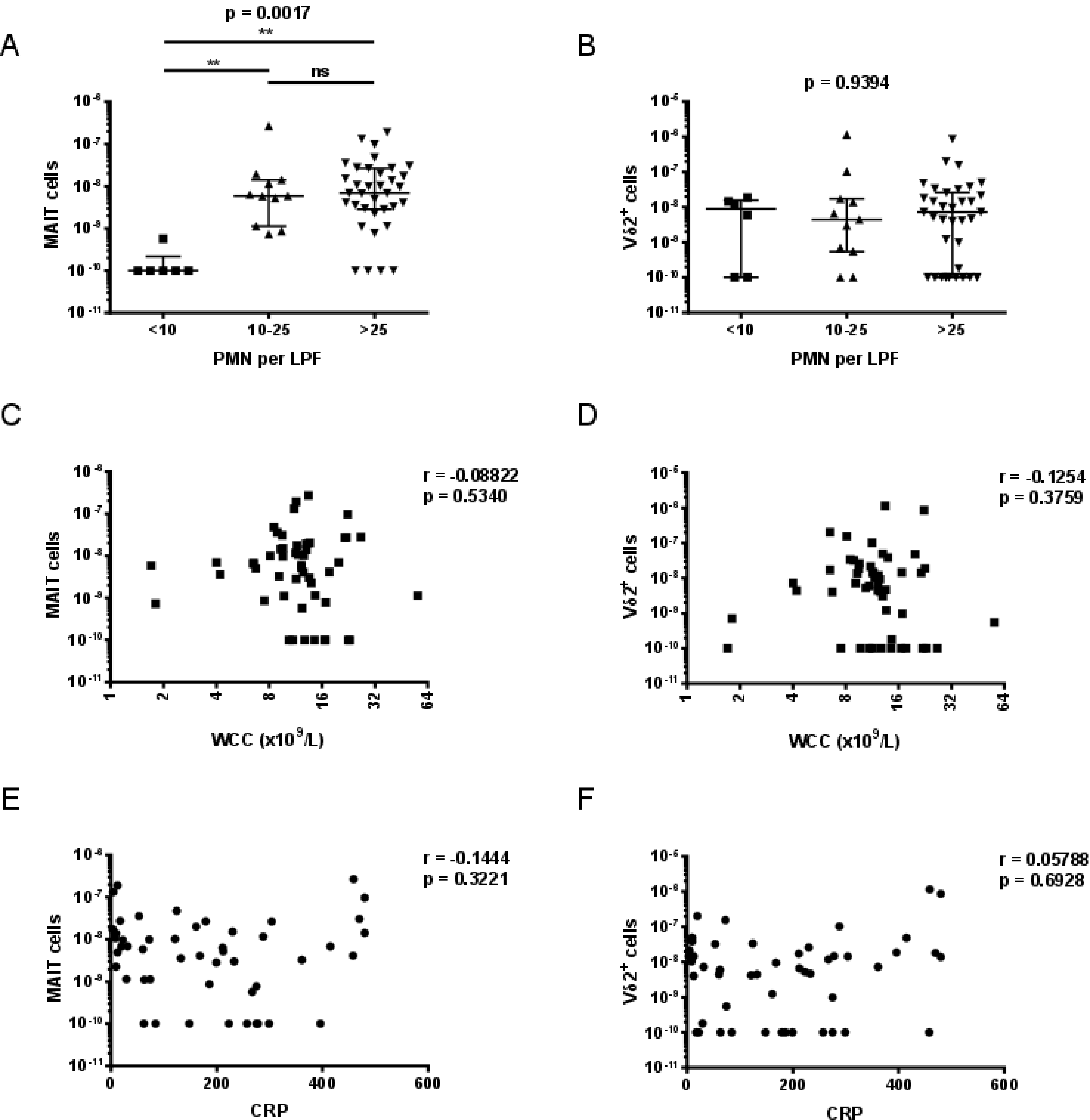

### Supplementary file 6

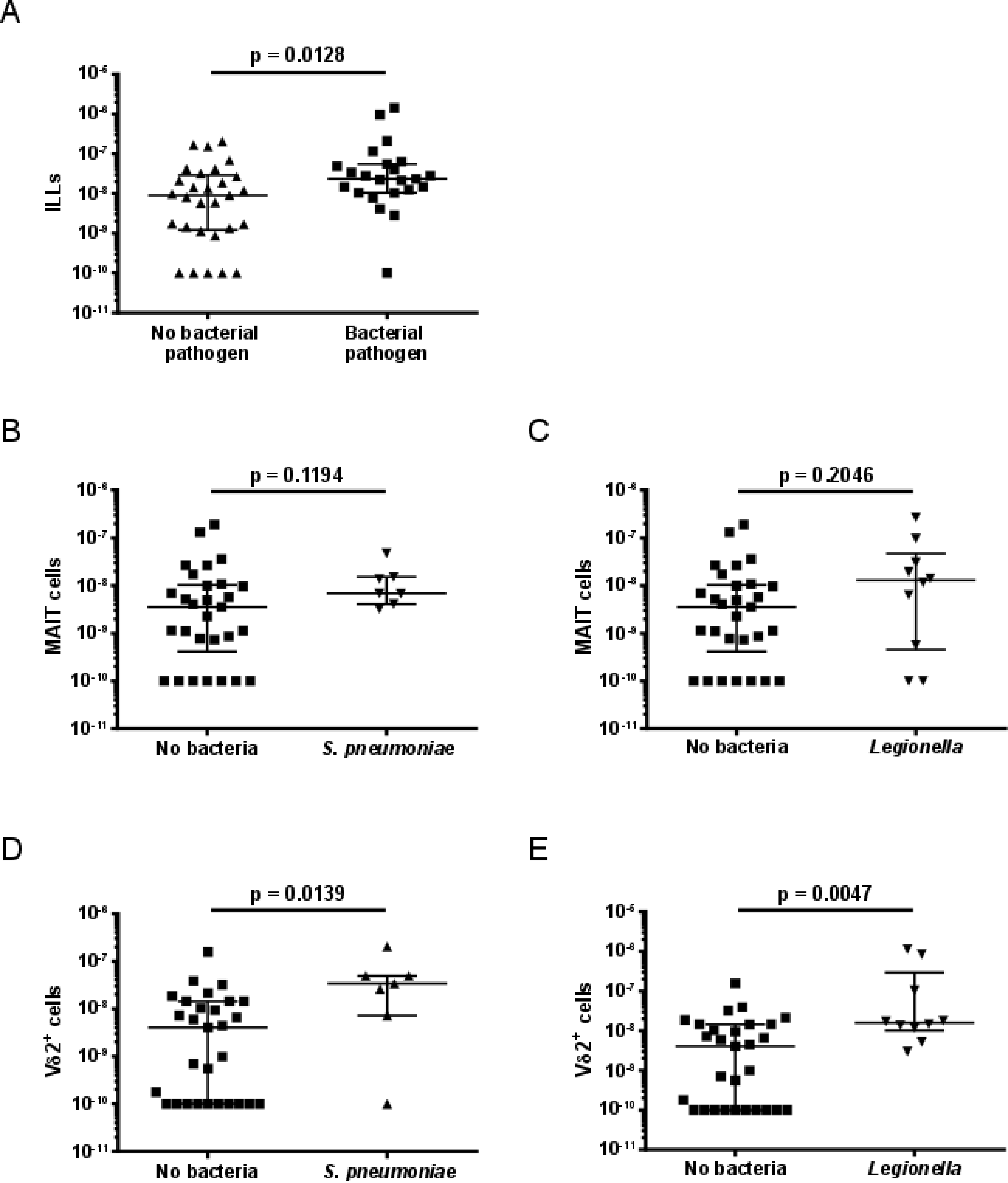

### Supplementary file 7

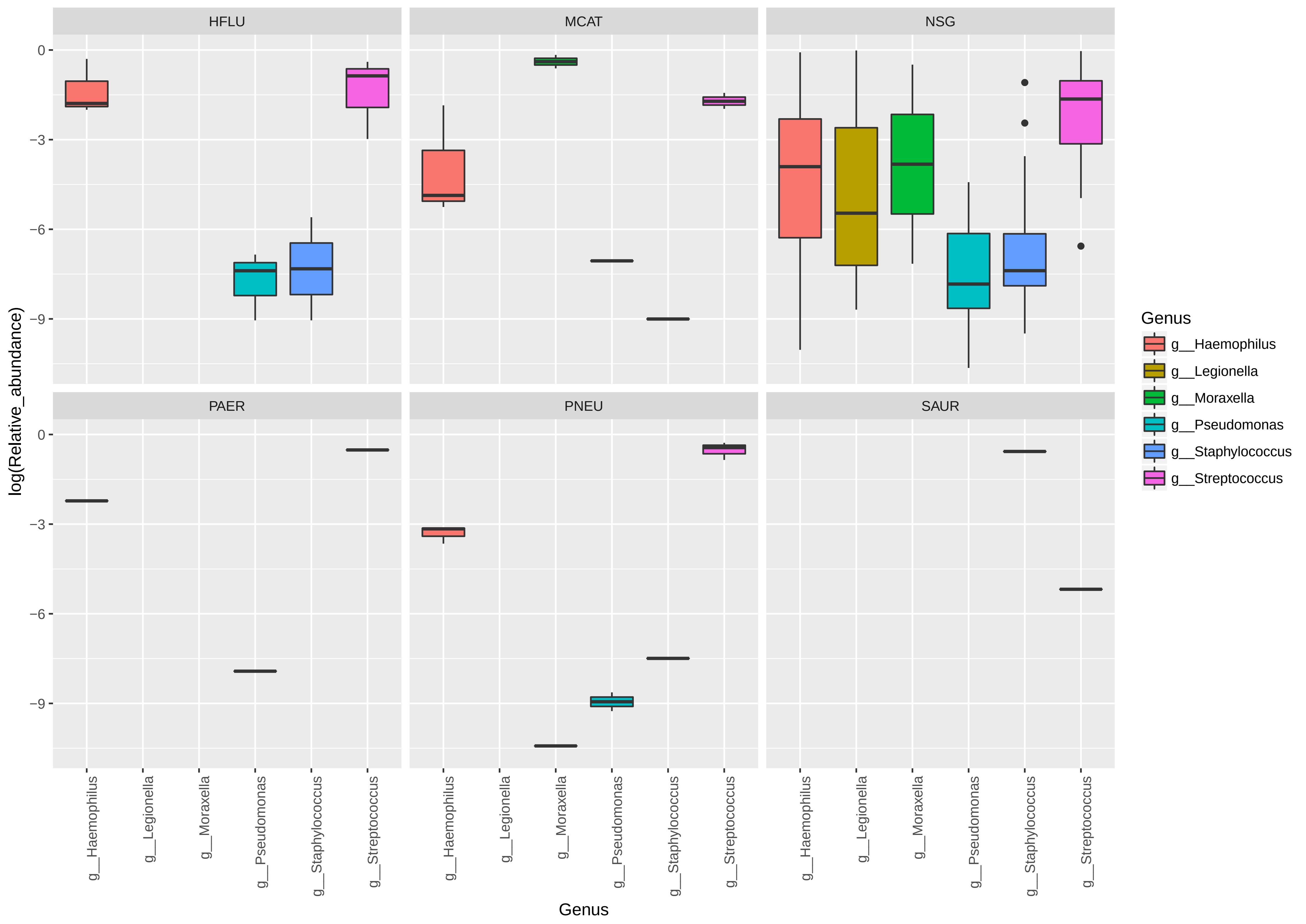

### Supplementary file 8

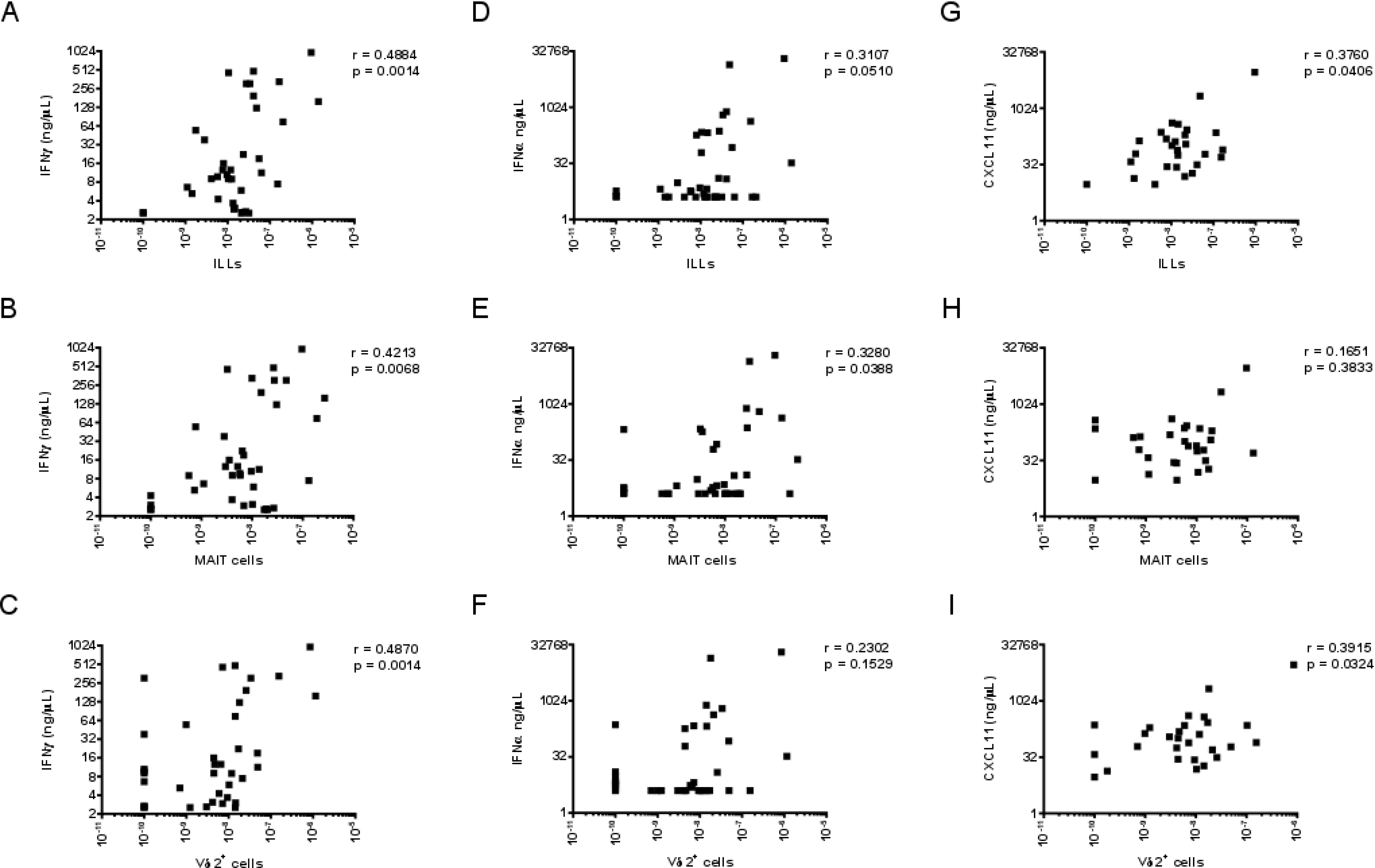
