## Supplementary material for "Role of Innate-like Lymphocytes in the Pathogenesis of Community Acquired Pneumonia"

**Supplementary Table 1. Primers and probes**

| Name | | Sequence (5’-3’) | Conc. (nM) | Reference |
| --- | --- | --- | --- | --- |
| hAV7S2 (FWD) | | TCCTTAGTCGGTCTAAAGGGTACAG | 400 | [51] |
| hAJ33 (REV) | | CCAGCGCCCCAGATTAA | 200 | [51] |
| TRAJ12 (REV) | | GTCCCACTCCCGAAGATCAATTT | 400 | This study |
| TRAJ20 (REV) | | TGTGGTTCCGGCTCCAAAG | 400 | This study |
| phAV7.2 (Probe) | | [6FAM]-TGAAAGACTCTGCCTCTTACCTCTGTGC-[BHQ1] | 400 | [51] |
| hVD2 (FWD) | | CTTGCACCATCAGAGAGAGATG | 400 | This study |
| hTRDJ1 (REV) | | CAGTCACACGGGTTCCTTT | 400 | This study |
| hTRDJ2 (REV) | | CTGGTTCCACGATGAGTTGT | 400 | This study |
| hTRDJ3 (REV) | | GGCTCCACGAAGAGTTTGAT | 400 | This study |
| hTRDJ4 (REV) | | GTACCTCCAGATAGGTTCCTTTG | 400 | This study |
| phVD2 (Probe) | | [56-FAM]-AGGGTCTTA-[ZEN]-CTACTGTGCCTGTGACA-[3IABkFQ] | 400 | This study |
| PAN23S-F (FWD) | | TCGCTCAACGGATAAAAG | 200 | [52] |
| PAN23S-R (REV) | | GATGAnCCGACATCGAGGTGC | 200 | [52] |
| β2M | **FWD**  **REV**  **Probe** | GTGCTGTCTCCATGTTTGATG  TCTGCTCCCCACCTCTAAG  [6-FAM]-AGGTTGCTC-[ZEN]-CACAGGTAGCTCTAGG-[IABkFQ] | 1X | [19] |
