## Supplementary material for "Role of Innate-like Lymphocytes in the Pathogenesis of Community Acquired Pneumonia"

**Supplementary Table 2:**

| **Process** | **KEGG genes** |
| --- | --- |
| Riboflavin biosynthesis | K01497, K14652, K01498, K11752, K00082, K20861, K20862, K21063, K21064, K02858, K14652, K00794, K00793, K00861, K20884, K11753, K00953 |
| C5 isoprenoid biosynthesis | K01662, K00099, K12506, K00919, K01770, K03526, K03527, K01823 |
