## Supplementary material for "Role of Innate-like Lymphocytes in the Pathogenesis of Community Acquired Pneumonia"

| | N (%) | ILL | | | | MAIT cells | | | | Vδ2+ T cells | | | |
|---|---|---|---|---|---|---|---|---|---|---|---|---|---|
| | | Median | 25th perc. | 75th perc. | P-value | Median | 25th perc. | 75th perc. | P-value | Median | 25th perc. | 75th perc. | P-value |
| Gender | | | | | | | | | | | | | |
| Males | 25 (48) | $2.22 \times 10^{-8}$ | $5.04 \times 10^{-9}$ | $3.73 \times 10^{-8}$ | 0.67 | $6.85 \times 10^{-9}$ | $1.00 \times 10^{-9}$ | $2.29 \times 10^{-8}$ | 0.29 | $6.60 \times 10^{-9}$ | $1.00 \times 10^{-10}$ | $1.58 \times 10^{-8}$ | 0.64 |
| Female | 27 (52) | $1.41 \times 10^{-8}$ | $1.76 \times 10^{-9}$ | $4.10 \times 10^{-8}$ | | $3.56 \times 10^{-9}$ | $7.71 \times 10^{-10}$ | $1.07 \times 10^{-8}$ | | $5.24 \times 10^{-9}$ | $7.05 \times 10^{-10}$ | $2.13 \times 10^{-8}$ | |
| COPD/Structural lung disease | | | | | | | | | | | | | |
| No | 33 (65) | $1.87 \times 10^{-8}$ | $4.94 \times 10^{-9}$ | $4.47 \times 10^{-8}$ | 0.32 | $5.88 \times 10^{-9}$ | $1.56 \times 10^{-9}$ | $2.35 \times 10^{-8}$ | 0.28 | $7.21 \times 10^{-9}$ | $5.45 \times 10^{-10}$ | $2.00 \times 10^{-8}$ | 0.26 |
| Yes | 18 (35) | $1.54 \times 10^{-8}$ | $1.41 \times 10^{-9}$ | $3.09 \times 10^{-8}$ | | $4.49 \times 10^{-9}$ | $5.74 \times 10^{-10}$ | $1.13 \times 10^{-8}$ | | $3.53 \times 10^{-9}$ | $1.00 \times 10^{-10}$ | $1.44 \times 10^{-8}$ | |
| Asthma | | | | | | | | | | | | | |
| No | 41 (79) | $1.87 \times 10^{-8}$ | $4.29 \times 10^{-9}$ | $4.82 \times 10^{-8}$ | 0.22 | $6.40 \times 10^{-9}$ | $9.96 \times 10^{-10}$ | $1.97 \times 10^{-8}$ | 0.53 | $7.24 \times 10^{-9}$ | $1.00 \times 10^{-10}$ | $2.68 \times 10^{-8}$ | 0.56 |
| Yes | 11 (21) | $7.69 \times 10^{-9}$ | $1.44 \times 10^{-9}$ | $3.18 \times 10^{-8}$ | | $4.11 \times 10^{-9}$ | $7.32 \times 10^{-10}$ | $1.51 \times 10^{-8}$ | | $4.70 \times 10^{-9}$ | $7.05 \times 10^{-10}$ | $1.44 \times 10^{-8}$ | |
| Duration of illness (days) | | | | | | | | | | | | | |
| 1-2 | 12 (24) | $2.23 \times 10^{-8}$ | $6.57 \times 10^{-9}$ | $2.79 \times 10^{-8}$ | 0.29 | $6.08 \times 10^{-9}$ | $1.42 \times 10^{-9}$ | $1.41 \times 10^{-8}$ | 0.41 | $7.17 \times 10^{-9}$ | $1.20 \times 10^{-10}$ | $1.83 \times 10^{-8}$ | 0.53 |
| 3-5 | 21 (41) | $1.18 \times 10^{-8}$ | $2.90 \times 10^{-9}$ | $4.10 \times 10^{-8}$ | | $4.05 \times 10^{-9}$ | $3.34 \times 10^{-10}$ | $1.34 \times 10^{-8}$ | | $6.60 \times 10^{-9}$ | $1.78 \times 10^{-9}$ | $1.45 \times 10^{-8}$ | |
| 6-8 | 4 (8) | $1.46 \times 10^{-9}$ | $1.00 \times 10^{-10}$ | $2.60 \times 10^{-8}$ | | $1.46 \times 10^{-9}$ | $1.00 \times 10^{-10}$ | $3.63 \times 10^{-8}$ | | $1.00 \times 10^{-10}$ | $1.00 \times 10^{-10}$ | $2.53 \times 10^{-8}$ | |
| >8 | 14 (27) | $1.79 \times 10^{-8}$ | $4.92 \times 10^{-9}$ | $8.93 \times 10^{-8}$ | | $8.27 \times 10^{-9}$ | $1.05 \times 10^{-9}$ | $2.40 \times 10^{-8}$ | | $6.61 \times 10^{-9}$ | $7.67 \times 10^{-10}$ | $3.64 \times 10^{-8}$ | |
| Antibiotics pre-admission | | | | | | | | | | | | | |
| No | 38 (74) | $1.38 \times 10^{-8}$ | $5.03 \times 10^{-9}$ | $3.55 \times 10^{-8}$ | 0.91 | $5.49 \times 10^{-9}$ | $1.13 \times 10^{-9}$ | $1.39 \times 10^{-8}$ | 0.58 | $5.92 \times 10^{-9}$ | $4.61 \times 10^{-10}$ | $2.05 \times 10^{-8}$ | 0.73 |
| Yes | 14 (27) | $2.12 \times 10^{-8}$ | $6.71 \times 10^{-10}$ | $8.93 \times 10^{-8}$ | | $8.73 \times 10^{-9}$ | $1.00 \times 10^{-10}$ | $3.18 \times 10^{-8}$ | | $8.14 \times 10^{-9}$ | $1.00 \times 10^{-10}$ | $1.89 \times 10^{-8}$ | |
| Sample type | | | | | | | | | | | | | |
| Expectorated | 43 (83) | $2.10 \times 10^{-8}$ | $5.76 \times 10^{-9}$ | $4.86 \times 10^{-8}$ | 0.06 | $6.76 \times 10^{-9}$ | $1.13 \times 10^{-9}$ | $1.92 \times 10^{-8}$ | 0.13 | $5.97 \times 10^{-9}$ | $1.79 \times 10^{-10}$ | $2.60 \times 10^{-8}$ | 0.20 |
| Induced | 9 (17) | $1.05 \times 10^{-8}$ | $4.81 \times 10^{-10}$ | $1.38 \times 10^{-8}$ | | $3.26 \times 10^{-9}$ | $4.16 \times 10^{-10}$ | $6.04 \times 10^{-9}$ | | $6.60 \times 10^{-9}$ | $1.00 \times 10^{-10}$ | $8.32 \times 10^{-9}$ | |
| Consolidation on chest x-ray | | | | | | | | | | | | | |
| Lobar | 31 (60) | $1.24 \times 10^{-8}$ | $2.82 \times 10^{-9}$ | $4.07 \times 10^{-8}$ | 0.76 | $4.93 \times 10^{-9}$ | $1.11 \times 10^{-9}$ | $1.16 \times 10^{-8}$ | 0.30 | $7.21 \times 10^{-9}$ | $1.79 \times 10^{-10}$ | $1.81 \times 10^{-8}$ | 0.61 |
| Multilobar | 21 (40) | $2.10 \times 10^{-8}$ | $2.94 \times 10^{-9}$ | $4.10 \times 10^{-8}$ | | $1.02 \times 10^{-8}$ | $8.16 \times 10^{-10}$ | $2.33 \times 10^{-8}$ | | $4.20 \times 10^{-9}$ | $1.00 \times 10^{-10}$ | $2.24 \times 10^{-8}$ | |
