## Supplementary material for "Role of Innate-like Lymphocytes in the Pathogenesis of Community Acquired Pneumonia"

| Bacterial species | Detection method | n |
|---|---|---|
| *Legionella longbeachae* | Sputum PCR only | 6 |
| | Sputum PCR + culture | 4 |
| *Streptococcus pneumoniae* | Sputum culture | 3 |
| | Blood culture | 1 |
| | Urine Antigen | 3[*][†] |
| *Haemophilus influenzae* | Sputum culture | 3[†] |
| *Moraxella catarrhalis* | Sputum culture | 3[*] |
| *Staphylococcus aureus* | Sputum culture | 1 |
| *Pseudomonas aeruginosa* | Sputum culture | 1 |

[*] Sputum culture grew *M. catarrhalis* and pneumococcal urinary antigen positive

[†] Sputum culture grew *H. influenzae* and pneumococcal urinary antigen positive
