## Supplementary material for "Role of Innate-like Lymphocytes in the Pathogenesis of Community Acquired Pneumonia"

| | MAIT | | | Vδ2+ T cells | | |
|---|---|---|---|---|---|---|
| | Spearman r | P value | Q value | Spearman r | P value | Q value |
| Riboflavin KEGGs (M00125) | | | | | | |
| K00794 | 0.15 | 0.3 | 0.5 | 0.03 | 0.84 | 0.98 |
| K00793 | 0.23 | 0.1 | 0.38 | 0.09 | 0.52 | 0.78 |
| K01498 | NA | NA | NA | NA | NA | NA |
| K01497 | -0.23 | 0.1 | 0.38 | -0.27 | 0.05 | 0.46 |
| K02858 | -0.07 | 0.61 | 0.68 | 0 | 0.98 | 0.98 |
| K11753 | 0.12 | 0.4 | 0.51 | 0.21 | 0.13 | 0.51 |
| K11752 | 0.02 | 0.89 | 0.89 | -0.15 | 0.27 | 0.51 |
| K14652 | 0.14 | 0.33 | 0.5 | 0 | 0.98 | 0.98 |
| K00861 | -0.21 | 0.13 | 0.38 | -0.15 | 0.28 | 0.51 |
| K00082 | -0.17 | 0.24 | 0.5 | -0.18 | 0.19 | 0.51 |
| Non-Mevalonate KEGGs (M00096) | | | | | | |
| K01823 | 0.12 | 0.38 | 0.44 | 0.09 | 0.54 | 0.62 |
| K01662 | -0.14 | 0.31 | 0.41 | -0.12 | 0.4 | 0.53 |
| K00099 | -0.25 | 0.08 | 0.25 | -0.24 | 0.08 | 0.2 |
| K03527 | -0.26 | 0.07 | 0.25 | -0.24 | 0.08 | 0.2 |
| K03526 | -0.24 | 0.09 | 0.25 | -0.23 | 0.1 | 0.2 |
| K12506 | -0.09 | 0.53 | 0.53 | -0.04 | 0.79 | 0.79 |
| K01770 | -0.21 | 0.14 | 0.26 | -0.19 | 0.17 | 0.28 |
| K00919 | -0.2 | 0.16 | 0.26 | -0.27 | 0.06 | 0.2 |

Q value = Benjamini and Hochberg’s FDR correction
